# pySuStaIn: a Python implementation of the Subtype and Stage Inference algorithm

**DOI:** 10.1101/2021.06.09.447713

**Authors:** Leon M. Aksman, Peter A. Wijeratne, Neil P. Oxtoby, Arman Eshaghi, Cameron Shand, Andre Altmann, Daniel C. Alexander, Alexandra L. Young

**Affiliations:** Stevens Neuroimaging and Informatics Institute, Keck School of Medicine, University of Southern California; Centre for Medical Image Computing, Departments of Computer Science and Medical Physics, University College London; Queen Square Multiple Sclerosis Centre, Department of Neuroinflammation, UCL Queen Square Institute of Neurology, Faculty of Brain Sciences, University College London; Department of Neuroimaging, Institute of Psychiatry, Psychology and Neuroscience, King’s College London

**Keywords:** Disease progression modeling, disease heterogeneity, disease subtyping, disease staging

## Abstract

Progressive disorders are highly heterogeneous. Symptom-based clinical classification of these disorders may not reflect the underlying pathobiology. Data-driven subtyping and staging of patients has the potential to disentangle the complex spatiotemporal patterns of disease progression. Tools that enable this are in high demand from clinical and treatment-development communities. Here we describe the pySuStaIn software package, a Python-based implementation of the Subtype and Stage Inference (SuStaIn) algorithm. SuStaIn unravels the complexity of heterogeneous diseases by inferring multiple disease progression patterns (*subtypes*) and individual severity (*stages*) from cross-sectional data. The primary aims of pySuStaIn are to enable widespread application and translation of SuStaIn via an accessible Python package that supports simple extension and generalization to novel modelling situations within a single, consistent architecture.

**Code metadata:** Current code version *v1*.*0*

Permanent link to code/repository used of this code version *https://github.com/ucl-pond/pySuStaIn*

Legal Code License *MIT*

Code versioning system used *git*

Software code languages, tools, and services used *Python*

Compilation requirements, operating environments & dependencies *Linux, Mac, Windows*

Support email for questions

## 1. Motivation and Significance

The Subtype and Stage Inference (SuStaIn) algorithm is a powerful tool for understanding the progression of heterogeneous diseases. SuStaIn uniquely identifies distinct disease progression patterns (*subtypes*) that account for temporal change. This is valuable because many progressive diseases are heterogeneous in nature and can naturally be described by a set of distinct subtypes [1], [2],[3], [4]. SuStaIn has been applied to a number of neurodegenerative diseases, including Alzheimer’s disease [5][6], frontotemporal dementia [5] and multiple sclerosis (MS) [7]. It has also been being applied to progressive lung disease [8].

In contrast to SuStaIn, most disease progression models try to find a single coherent picture of how a disease evolves from early to late stages based on cross-sectional or short-term longitudinal snapshots of disease progression within individuals. Such models assume that all individuals follow the same pattern of progression [9], [10], [11], [12]. SuStaIn generalizes these models to infer multiple patterns of progression (from equivalent data). It does so via a spatiotemporal clustering approach that disentangles disease subtypes (i.e. distinct spatial patterns of progression) from disease stages (i.e. severity or the degree of temporal progression within a particular subtype). Importantly, the subtypes and stages inferred by SuStaIn cannot be resolved by clustering directly on subtype, which does not account for heterogeneity in disease severity within-cluster, or on stage, which does not account for subtype heterogeneity within-cluster.

The motivation for developing the pySuStaIn package is to expand the accessibility of the algorithm via an open-source Python implementation (originally MATLAB). This empowers more users to try the algorithm on their data sets and enables more developers to easily extend the code or compare it with other methods. Another major motivation is to increase the flexibility of the algorithm to handle multiple disease progression models. The original SuStaIn implementation was based on a linear z-score based likelihood function; pySuStaIn generalizes SuStaIn to handle arbitrarily defined data likelihood terms as derived classes within an object-oriented architecture. This allows direct plug-in of new models. As an initial demonstration, we implemented three models as derived classes within this framework: (i) the original continuous model of disease progression relative to a control population using the z-score based likelihood (*ZScoreSustain*); (ii) the ‘normal-versus-abnormal’ model which uses a data likelihood based on a mixture of normal and abnormal distributions, as in the Event Based Model (*MixtureSustain*; [9], [13], [Firth-2020]); and (iii) a model for discrete ordinal data, that can be used for biomarkers based on visual ratings, neuropathological ratings or certain cognitive tests (*OrdinalSustain*).

The pySuStaIn package is intended to be flexible and easy to use: the user chooses the type of likelihood and sets a few parameters controlling the number of subtypes to be inferred, the number of Markov chain Monte Carlo (MCMC) iterations and expectation maximization (EM) start-points, and whether to use parallelization. It is also intended to be easy to extend: new disease progression models can be added as implementations of *AbstractSustain* with an appropriately defined likelihood. Simulation code and Jupyter notebooks are provided to help users understand these functionalities.

## 2. Software Description

pySuStaIn is written in Python 3 and uses the NumPy and SciPy numerical packages. It uses the Pathos package for parallelizing the start-points of several EM-based computations described later. We used Pathos instead of Python’s default multiprocessing package as it allows separate random seeds across processes, which is important because pySuStaIn makes extensive use of randomly permuted sequences. The following sections describe pySuStaIn’s software architecture and its major functionalities. Several code snippets are included to aid understanding.

### 2.1 Software Architecture

The SuStaIn algorithm was described in detail by Young et al. [5]. Briefly, SuStaIn infers increasingly complex models of disease progression using a set of cross-sectional training samples (a subjects-by-features matrix). Ideally, these samples should adequately capture the dynamics of some underlying heterogeneous disease, in terms of both the variability of progression and disease severity. Given such data, SuStaIn begins by inferring a single sequence of events that characterizes disease progression from early to late stages. Each successive iteration increases model complexity by adding a subtype, up to a specified maximum. The original implementation [5] used a z-score based data likelihood to find the maximum likelihood event sequence for each subtype. Each biomarker was associated with a fixed set of z-score based events; an event corresponded to a biomarker value exceeding (for example) one, two or three standard deviations relative to a control population mean. pySuStaIn generalizes the algorithm to accept customizable data likelihoods, e.g., based on a mixture model. The user is free to choose the data likelihood that best fits their problem from among the available implementations (Figure 1), or to contribute their own.

**Figure 1.**
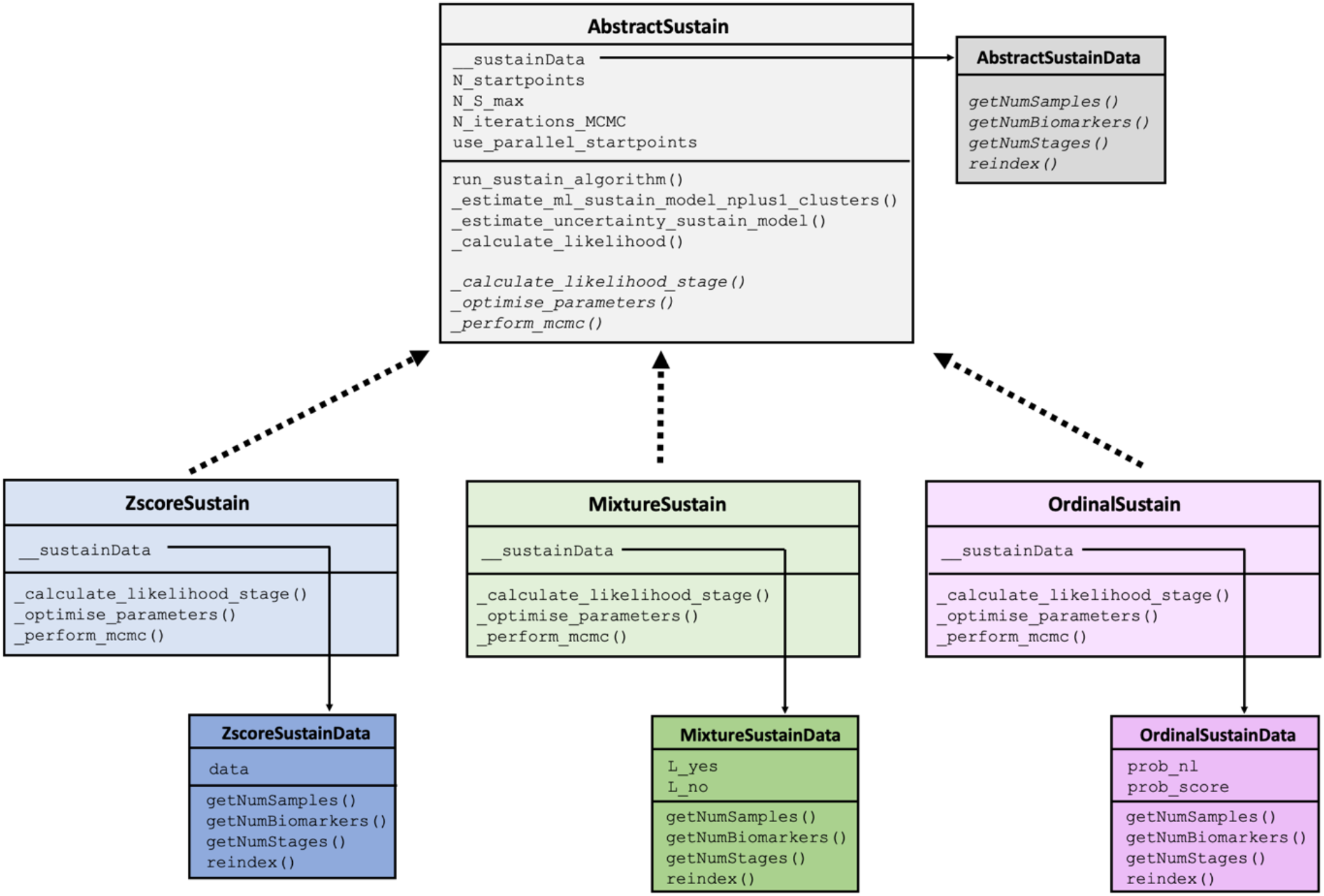
The architecture of pySuStaIn.

The core functionality of the algorithm is implemented within the *AbstractSustain* class, which, as its name suggests, is an abstract class that cannot be instantiated directly. As shown in Figure 1, each child class then implements its own `_calculate_likelihood_stage()`, _`optimise_parameters()` and _`perform_mcmc()` methods, all of which depend on the child class’ particular data likelihood function. Each child class also has a `sustainData` member variable which is an instance of an *AbstactSustainData*-derived class. In the case of the *ZscoreSustain* class, this is a *ZscoreSustainData* object with an internal data variable containing z-scored biomarkers for all subjects, where z-scoring is done externally. For *MixtureSustain*, this is a *MixtureSustainData* object with internal `L_yes` and `L_no` matrices storing the probabilities that a biomarker measure belongs, respectively, to the patient or control distributions of a mixture model. Storing the probabilities rather than the data itself allows for complete flexibility in the form of the probability distribution used to model the probability an event has or hasn’t occurred. These matrices are typically generated via a mixture modelling procedure that can be either performed in pySuStaIn or done externally. In the example code provided (`simrun.py`), mixture models are built for each biomarker using either Gaussian mixture modelling (setting `sustainType` to `mixture_gmm`) or kernel density estimation mixture modelling (setting `sustainType` to `mixture_kde`) [14]. *OrdinalSustain* similarly stores an internal *OrdinalSustainData* object with `prob_nl` and `prob_score` matrices respectively storing the probabilities of each biomarker being ‘normal’ (similar to the expected score in a control population) or having each particular score. As with *MixtureSustain*, this formulation allows complete flexibility for the user to choose the form of the distributions.

To aid in understanding our implementation, we briefly explain some of the most important methods within *AbstractSustain*. The `_estimate_ml_sustain_model_nplus1_clusters()` method within `run_sustain_algorithm()`, which starts the algorithm after initialization, is responsible for inferring the desired number of subtypes. It is based on the principle that splitting an existing subtype into two subtypes is computationally much simpler than splitting a whole dataset into multiple subtypes. Initially, the `_find_ml()` function uses greedy expectation maximization (EM) to find a maximum likelihood based biomarker sequence that describes all subjects’ progression (i.e. a one-subtype model). Once this sequence is inferred, two separate methods generalize the algorithm to multiple subtypes. The first is `_find_ml_split()`, depicted in Figure 2a, which finds the best split of an existing event sequence into two subtypes. Within this method, the `n` subjects described by a sequence are randomly split into two subgroups and greedy EM is run separately for each subgroup to find an optimal sequence describing that subgroup’s progression. The second is `_find_ml_mixture()`, which uses the newly split sequence and the unsplit sequences from the previous iteration as a starting point to optimise the full set of sequences. For example, as depicted in Figure 2b, if SuStaIn is trying to infer a three-subtype model based on two previously inferred sequences S_1_ and S_2_, it will try two different splits and evaluate how well each of them fits the data. It will split S_1_ into S_1,1_ and S_1,2_ and use greedy EM to find maximum likelihood sequences starting with S_1,1_, S_1,2_ and S_2_. Similarly, it will split S_2_ into S_2,1_ and S_2,2_ and do the same with S_1_, S_2,1_ and S_2,2_. Using maximum likelihood it will choose which of these two sets of optimized sequences best describes the whole training dataset and choose that set as the new three-subtype model. To find a four-subtype model the algorithm then splits each of the three subtypes and finds which of the three resulting four-subtype models best describes the data and so on.

**Figure 2.**
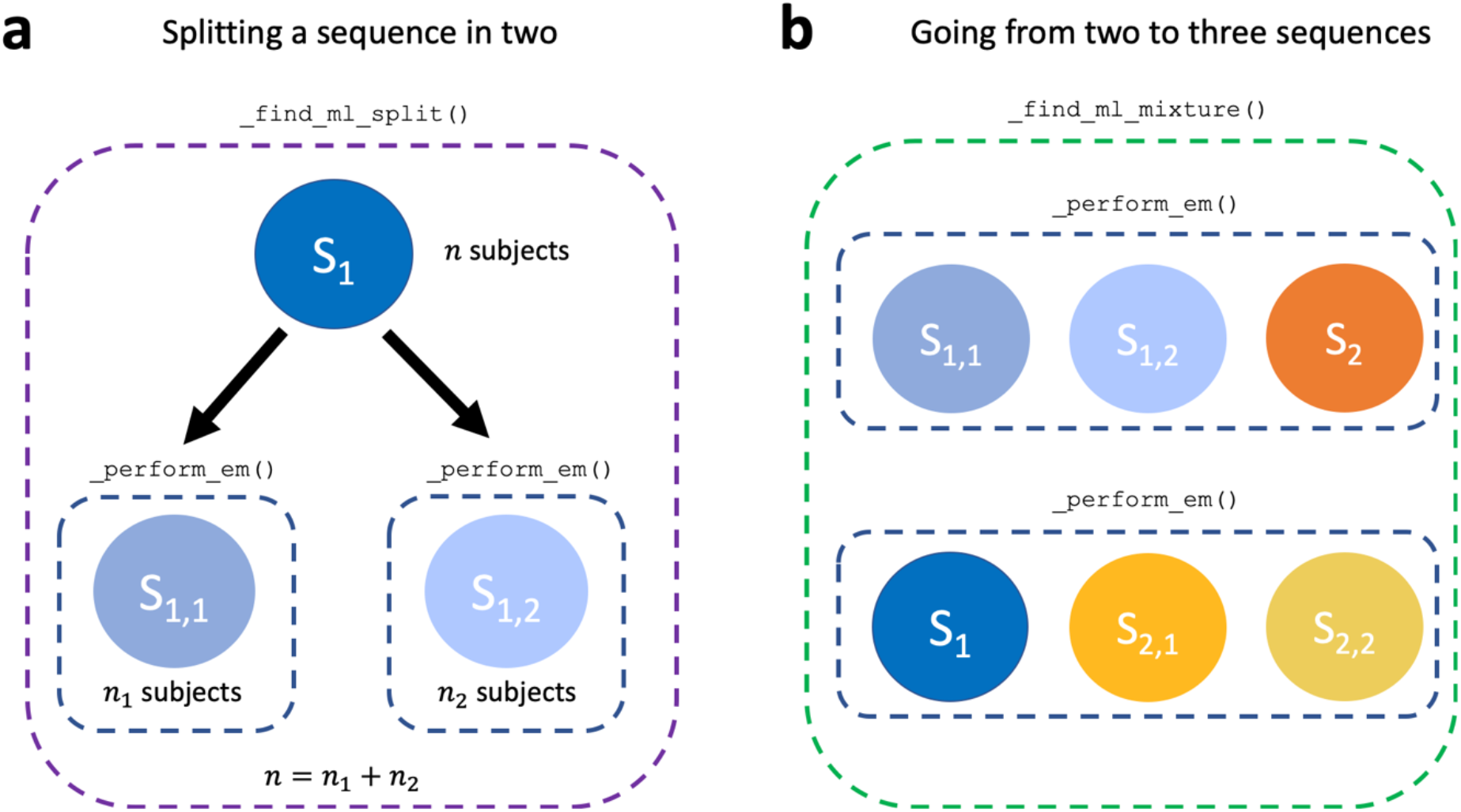
Core computations within the `_estimate_ml_sustain_model_nplus1_clusters()` method of *AbstractSustain*.

Once the sequences have been inferred using the above procedure, `_estimate_uncertainty_sustain_model()` uses MCMC to estimate the positional uncertainties for each sequence. Each sequence is permuted by swapping two randomly chosen places and the likelihood of the data under the permutation is evaluated. This procedure is performed a set number of times (typically one hundred thousand or one million). From this a positional variance diagram (PVD) can be built to visualize how often each biomarker appears in each position (see [9] for further explanation).

### 2.2 Software Functionalities

The major functionalities of pySuStaIn are: (i) flexible choice of data likelihood; (ii) data preparation, which depends on the likelihood; (iii) SuStaIn-based inference to find biomarker progression sequences for the specified number of subtypes; (iv) visualizations of the inferred sequences; (v) estimation of the most likely subtype and stage of each subject based on the inferred model; and (vi) tools to aid in model selection. These functionalities are depicted in Figure 3 and described in greater detail in the following subsections.

**Figure 3.**
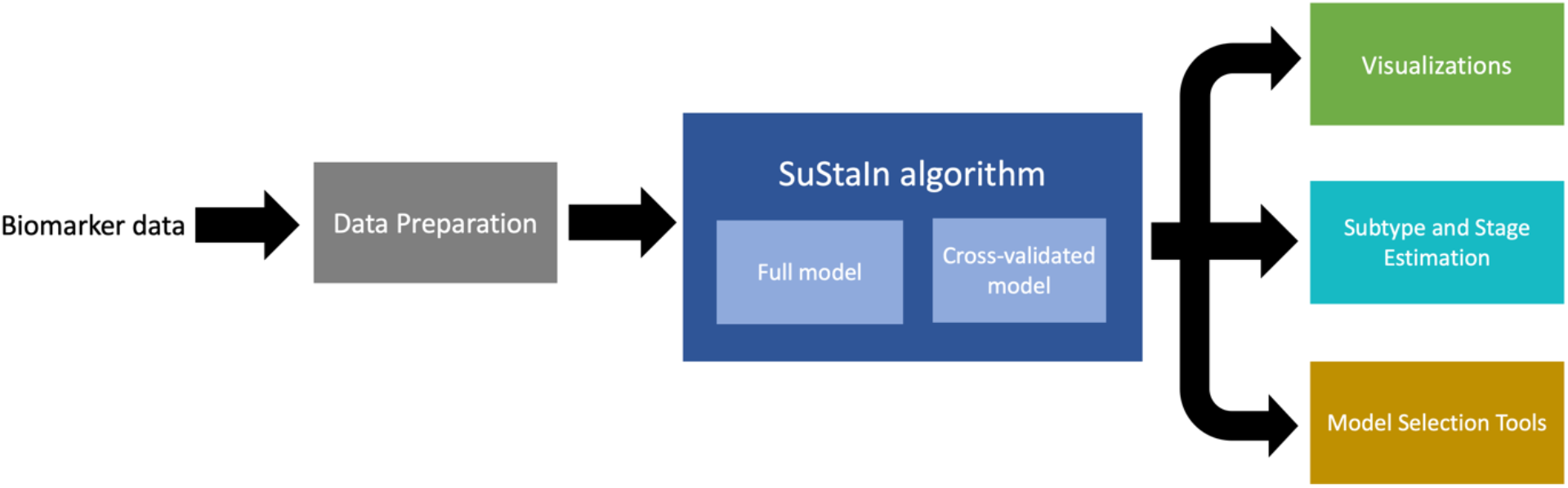
Depiction of the major functionalities of pySuStaIn, as a sequence of operations that begins with an input biomarker data matrix, followed by a data preparation step that depends on the chosen data likelihood, then the SuStaIn algorithm run on both full and cross-validated data and finally a set of outputs consisting of: (i) visualizations of the inferred models; (ii) estimates of the most likely subtype and stage for training and test subjects; and (iii) a set of model selection tools.

#### 2.2.1 Flexible choice of data likelihood

One of the most important functionalities of pySuStaIn is that it allows users to choose among the existing implementations of *AbstractSustain* and to easily add new ones. Each represents a different underlying disease progression model, defined by a unique data likelihood. For example, *ZscoreSustain* represents continuous accumulation of abnormality relative to a control population using a z-score based likelihood while *MixtureSustain* models transitions from normal to abnormal measures using a mixture model based likelihood. *OrdinalSustain* models transitions from one score to another using a categorical likelihood. Other disease progression models can easily be added by deriving from *AbstractSustain* and *AbstractSustainData*. Importantly, in all such derived classes the core algorithm is unchanged: it always uses greedy EM and MCMC to infer sequences of events (however defined) that best explain the available data.

#### 2.2.2 Data preparation

Data preparation varies depending on which data likelihood is used. In the case of the z-score likelihood, the *ZscoreSustain* class assumes that users will z-score the input data themselves and hence there is no data preparation in this case. For clarity, z-scoring should be done with respect to a control population, so that z-score based events can be interpreted as departures from normality. In `simrun.py` this functionality is showcased when `sustainType` is set to `zscore`.

In the case of the mixture model likelihood, as mentioned above, *MixtureSustain*’s constructor expects `L_yes` and `L_no` likelihood matrices. While users are free to build their own mixture models to generate these matrices, pySuStaIn implements Gaussian mixture models (GMMs) and kernel density estimation based mixture models (KDE-MMs), both of which have been previously used with the Event Based Model [9], [13], [14]. Within `simrun.py`, simulated subjects assigned earliest stages are used as controls and those in latest stages as cases. Mixture models are then fit using either `fit_all_gmm_models` or `fit_all_kde_models`, depending on whether `sustainType` is set to `mixture_gmm` or `mixture_kde`, respectively. As a general rule, GMMs should be used when the normal and abnormal distributions for each biomarker are suspected to be normally distributed while KDE-MMs should be used in more general cases when these values are not necessarily Gaussian, e.g., they are heavy-tailed or asymmetric.

#### 2.2.3 SuStaIn-based inference

Once data has been prepared, pySuStaIn is ready to run the SuStaIn algorithm to infer the specified number of subtypes. After an *AbstractSustain* type object (specifically, a *ZscoreSustain, MixtureSustain* or *OrdinalSustain* object) is initialized and `run_sustain_algorithm()` is called, SuStaIn proceeds to infer iteratively more complex models, beginning with a one-subtype model describing all subjects’ progression and ending with an `N_S_max`-subtype model, with `N_S_max` passed in on initialization. As SuStaIn can be computationally demanding, particularly with a large sample size, a large number of biomarkers (especially if the z-score likelihood is used), and/or a high `N_S_max`, pickle files are used to save the progress of the algorithm at each iteration of this procedure. Pickle files are saved within the /`pickle_files` subfolder of the `output_folder` directory, which is also passed in on instantiation. This allows the program to be restarted so that if, for example, the program has been previously run with `N_S_max` set to two and is subsequently set to three. In this case the algorithm does not have to recompute the one-subtype and two-subtype models to find a three-subtype model.

SuStaIn can also be run within a cross-validation scheme via the `cross_validate_sustain_model()` function, which accepts a list of test sample indices for each fold. The training indices for a fold are the set difference between all training samples and the given test samples. SuStaIn is then run on the fold’s training data, up to the specified number of subtypes, with each model saved as a fold-specific pickle file.

#### 2.2.4 Visualization of inferred subtypes

Once a SuStaIn model has been fit, positional variance diagrams (PVDs) are an intuitive way of visualizing each subtype’s inferred event ordering while accounting for the MCMC-based positional uncertainty estimates. For example, Figure 4a depicts a PVD showing the true sequences from a three-subtype model used to generate simulated data. Each biomarker has three well-defined z-score based events, with no uncertainty in the ordering. Figure 4b on the other hand, depicts a PVD of SuStaIn’s inferred sequences, where the uncertainty in the positioning of certain events is clearly seen.

**Figure 4.**
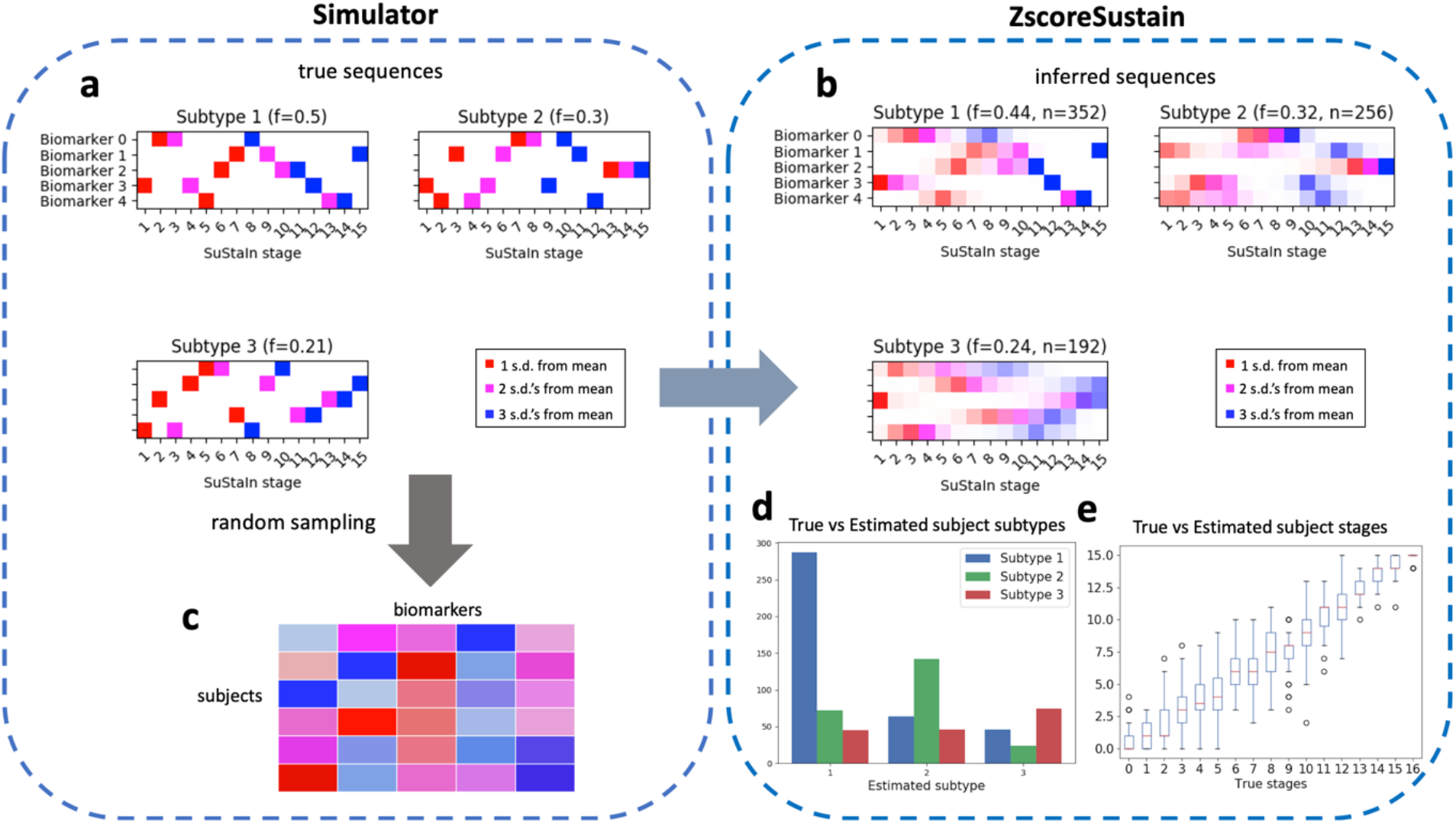
Schematic of simulated z-scored data as implemented in `simrun.py`. Inference of three subtypes via *ZScoreSustain* based on the simulated input data, with: (plot **a)** true event sequences for each subtype; and (**b)** sequences inferred by SuStaIn given a (**c**) subjects × features input matrix of z-scores for subjects with randomly-sampled subtypes and stages. A comparison of estimated versus true subtypes and stages are depicted in (**d**) and (**e**), respectively.

pySuStaIn generates PVDs for both full models (i.e. trained on all samples) and cross-validated models. Cross-validated PVDs are formed by finding the best one-to-one match between each fold’s and the full model’s sequences (based on Kendall’s tau correlation measure) so that subtypes’ MCMC samples can be stacked across folds. This is done because changing the training data (as in cross-validation) can change the ordering of events within inferred subtypes. Figures 4b and 4c depicts PVDs from the full and cross-validated models respectively, showing the additional uncertainty in the cross-validated version, as expected.

#### 2.2.5 Subtype and stage estimation

Given a SuStaIn model, consisting of the maximum likelihood and MCMC sequences, the `subtype_and_stage_individuals()` function estimates a most likely subtype and stage for each training sample. This is done by first calculating a probability distribution over all of the possible subtypes and stages for every subject based on the model and the given subject’s biomarker values. To find the most likely subtype we sum over all possible stages, choosing the subtype with the highest marginal probability. Similarly, to find the most likely stage, we sum over all subtypes, choosing the most probable stage for every subject. Figures 4d and 4e depict true versus estimated subtypes and stages for the inferred model built on simulated z-score data in `simrun.py`. The simulation writes the estimated subtype and stages to the `Subject_subtype_stage_estimates.csv` file. pySuStaIn can also estimate the subtypes and stages for unseen test samples via the `subtype_and_stage_individuals_newData()` function, which expects data in the same format as the training data (z-scored in *ZscoreSustain*; `L_yes`, `L_no` in *MixtureSustain*; `prob_nl`, `prob_score` in *OrdinalSustain*).

Note that users are free to derive alternative subtype and stage assignments using the three-way `prob_subtype_stage` matrix returned by this function (see Section 2.3).

#### 2.2.6 Tools for model selection

SuStaIn infers models of increasing complexity up to a maximum number of subtypes specified by `N_S_max`. Users must use their own discretion to select the most appropriate model. In general, models with more subtypes will better describe the training data (at the risk of overfitting), but will be harder to interpret. pySuStaIn provides several tools to aid model selection: (i) an MCMC likelihoods figure (`MCMC_likelihoods.png`), to help in visually comparing models’ in-sample model fits (see Figure 5f); (ii) after cross-validation is finished, it prints to the terminal the cross-validation information criterion (CVIC) for one-subtype to `N_S_max`-subtype models [5]; and (iii) out-of-sample log likelihoods, averaged across MCMC samples, for each cross-validation fold as both a figure (`Log_likelihoods_cv_fold.png`; Figure 5g) and a file (`Log_likelihoods_cv_fold.csv`).

**Figure 5.**
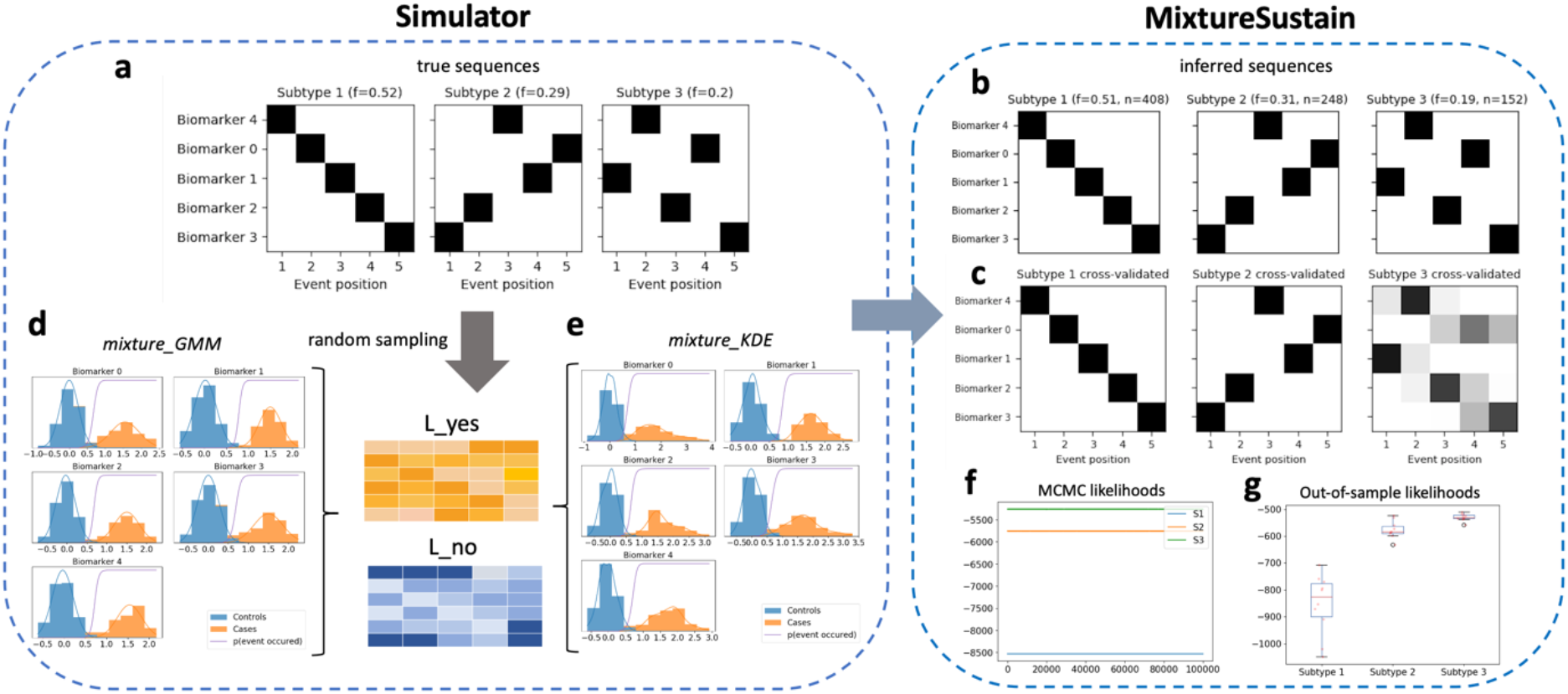
Schematic of simulated mixture model data as implemented in `simrun.py`, with the two available mixture model types shown (`mixture_GMM` or `mixture_KDE`, based on Gaussian mixture modelling or kernel density estimation, respectively). Inference of three subtypes using *MixtureSustain* on the simulated input data (`L_yes` and `L_no` matrices). Each matrix is a subjects × features matrix storing the probability of subjects’ observations belonging to the mixture-model-derived case (`L_yes`) or control (`L_no`) distribution. Figures shown are from the `mixture_GMM` style; those from the `mixture_KDE` style are very similar.

It is generally recommended to select models using either the CVIC or out-of-sample log likelihoods (either visually and/or via statistical tests between models of increasing complexity) than via in-sample MCMC likelihoods, as cross-validation approximates the generalizability of models to unseen data.

### 2.3 Sample code snippets

The following code snippets show how to initialize a *ZscoreSustain* object and run SuStaIn.

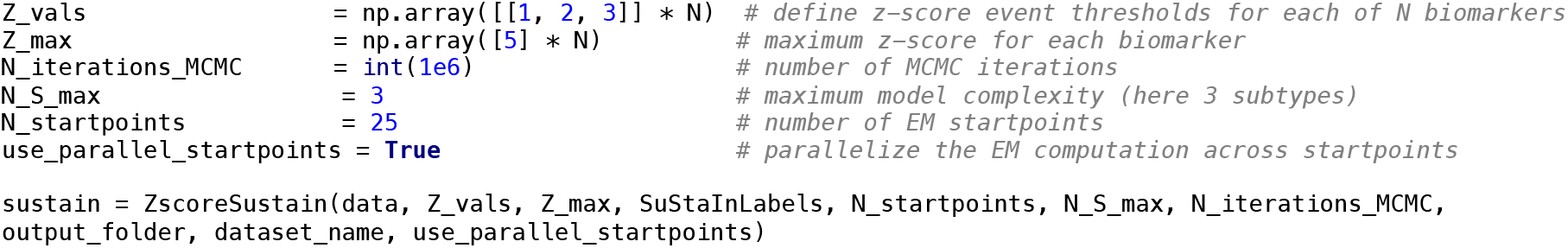

where `data` is a matrix of size `M` × `N`, where `M` is the number of training samples and `N` is the number of features and `SuStaInLabels` is a list of biomarker names for plotting. `Z_vals` specifies the z-score event thresholds for each biomarker as a matrix. In the above example, each of the `N` biomarkers is assigned three z-scores (1, 2 and 3). Users can also assign different numbers and values of z-scores each biomarker by setting some values in the `z_vals` to zero. For example, the following code:

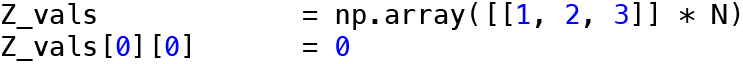

sets the first biomarker’s z-scores to 0, 2 and 3 and the rest of the biomarkers’ z-scores to 1, 2, 3 as before. If this version of `Z_vals` is passed to the *ZScoreSustain* constructor, the first biomarker will only have two associated z-score thresholds (2 and 3) and overall there will be `3N-1` stages in the inferred sequences rather than `3N`. By setting some elements of `Z_vals` to zero in this way, the user can fully customize the set of z-scores for each biomarker.

Once the `sustain` object has been initialized SuStaIn can be run via:

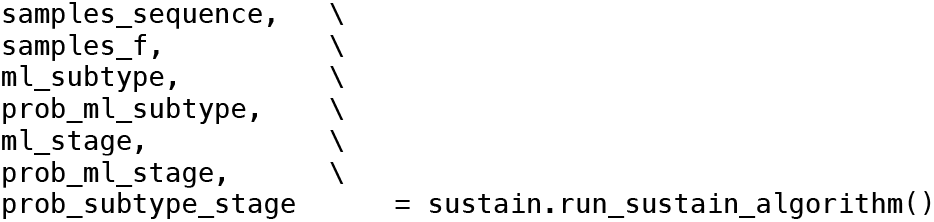

where `sample_sequences` and `samples_f` are the event sequences and fractions of subjects per subtype across MCMC samples, `ml_subtype` and `prob_ml_subtype` are the maximum likelihood subtype and associated subtype probability per sample, and `ml_stage` and `prob_ml_stage` are maximum likelihood stage and stage probability per sample. The `M` × `N_stages` × `N_S_max` matrix `prob_subtype_stage` stores the full probability distributions over all possible subtypes and stages for every sample, from which we derive the `prob_ml_subtype` and `prob_ml_stage` vectors.

Additionally, an instructional Jupyter notebook is also available in the `/notebooks` subdirectory.

## 3. Illustrative Examples

Illustrative examples of pySuStaIn are shown in Figures 4 and 5 for both the z-score and mixture likelihood styles of SuStaIn, which are the most commonly used implementations at present. These are produced by the `simrun.py` simulation code available in the `/sim` subdirectory. The simulator proceeds by randomly sampling a subtype (between zero and `N_S_ground_truth`, the number of ground truth subtypes, set to three in both cases) and stage (between zero and the total number of stages) for a set of 800 subjects. Ground truth subtypes are sampled from a discrete distribution with probabilities of 0.5, 0.3 and 0.2 assigned to the three subtypes. Ground truth stages are sampled from a uniform distribution.

Using randomly generated ground truth sequences (shown in Figures 4a and 5a) and these subtype and stage assignments, a training dataset is generated for input to the SuStaIn algorithm. The maximum number of subtypes to infer, `N_S_max`, is also set to three. Figures 4b and 5b show the inferred sequences for the three subtype models. In both cases there is a close correspondence to the true sequences, showing the ability of SuStaIn to recover the true underlying patterns of biomarker progression in a purely data-driven manner. Importantly, SuStaIn has a built-in quantification of uncertainty (positional variance, evident when comparing the inferred sequence in 4b to ground truth in 4a) which realistically reflects under-sampling across stages. Figures 4d and 4e depict the correspondence between the true (randomly generated) subtypes and stages of the training subjects and their estimated counterparts for the z-score likelihood simulation. Figures 5f and 5g depict the in-sample (MCMC likelihoods) and out-of-sample (cross-validation fold likelihoods) for the mixture simulation, showing that, as expected, the three-subtype model better explains the training data than simpler models.

## 4. Impact

Chronic diseases present enormous personal and societal challenges that will likely increase with the aging of worldwide populations. These include neurodegenerative diseases such as Alzheimer’s, Parkinson’s, and MS, as well as respiratory diseases such as chronic obstructive pulmonary disease (COPD). Understanding such complex multifactorial diseases necessarily involves accounting for heterogeneity using methods that group subjects with similar spatiotemporal progression patterns. SuStaIn is unique in its ability to find such subtypes in an objective, data-driven manner, using cross-sectional information from a suitably diverse set of samples.

The pySuStaIn package is intended to make SuStaIn widely accessible, easy to use and applicable to different modelling scenarios. It is already having a significant impact on the basic understanding of a variety of neurodegenerative diseases [15] and, in the longer term, will impact both clinical trials and clinical practice. pySuStaIn’s key features are: (i) it simplifies the process of implementing new subtyping models; (ii) it broadens the reach of the algorithm through Python; (iii) it is substantially faster than the original MATLAB implementation, due to several code optimizations, enabling more complex models to be fit; (iv) it has parallelized EM start-points for additional speed; (v) it integrates previously disparate disease progression models into a single package; and (vi) it adds both simulations and notebooks to make SuStaIn easier to understand and use.

Four recent studies illustrate how pySuStaIn has already enabled researchers to better understand both neurodegenerative and lung diseases:

- It has been used to characterize the spatiotemporal spread of Alzheimer’s related tau protein pathology throughout the brain. This is the largest study using tau PET imaging to date, analyzing over 1100 individuals from across five studies, with the aim of capturing as much tau heterogeneity as possible. It found four distinct subtypes, two of these were consistent with previous studies and two were novel subtypes which resembled atypical variants of Alzheimer’s. This study used the z-score likelihood, so that the interpretation of abnormality was relative to cognitively normal individuals [6].
- A related study investigated the spread of Alzheimer’s related amyloid and tau protein pathologies using PET imaging in 400 individuals from a single study. This study found that subjects fell into two basic subtypes: an amyloid-first subtype, in which amyloid pathology first appears in the brain, and a tau-first subtype. This model favors the dual pathway hypothesis of Alzheimer’s progression over the amyloid cascade hypothesis, the prevailing amyloid-centric model. This study used the mixture model likelihood, interpreting disease stages as transitions from distinctly normal to distinctly abnormal measurements [16].
- SuStaIn has also been used to find MRI-based subtypes of MS, showing that there are three distinct MRI-driven subtypes are better associated with disability progression than the current subtyping system that is based on four symptom-based subtypes. Importantly, the study also showed that only one of the identified subtypes showed a significant treatment response in randomised controlled trials. This study used the z-score likelihood [7].
- SuStaIn identified two major patterns of lung damage in Chronic Obstructive Pulmonary Disease (COPD) [8]: ‘tissue-airway’, which affected lung tissue early on, and ‘airway-tissue’, which affected the lung airways first. These patterns could be used to identify otherwise healthy smokers at risk of COPD at follow-up, suggesting that SuStaIn can be used for very early stratification in COPD. To date pySuStaIn has been developed and used by researchers at UCL, along with close collaborators. However, the intended user group for this package is the wider community of researchers and clinicians who are focussed on understanding neurodegenerative and other progressive diseases. We anticipate that pySuStaIn will facilitate this, as has been the case with event-based model code that is now finding broader use among the community [17].

## 5. Conclusions

We have presented pySuStaIn, a python-based implementation of the SuStaIn algorithm, a paradigm-shifting approach to understanding heterogeneity within progressive processes such as chronic diseases. pySuStaIn aims to widen the accessibility of this algorithm via an open source Python implementation. Our object-oriented implementation enables user-defined sub-models, which can be easily added to in the future, increasing the applicability of the algorithm. Inclusivity and accessibility is enhanced by providing code examples, visualizations to aid model interpretation, and tools for model selection.

## Conflict of Interest

We wish to confirm that there are no known conflicts of interest associated with this publication and there has been no significant financial support for this work that could have influenced its outcome.

## Acknowledgements

This project has received funding from the European Union’s Horizon 2020 research and innovation program under grant agreement No. 666992. PAW was supported by a MRC Skills Development Fellowship (MR/T027770/1). NPO is a UKRI Future Leaders Fellow (MR/S03546X/1). AA holds an MRC eMedLab Medical Bioinformatics Career Development Fellowship. NPO and DCA acknowledge funding from the National Institute for Health Research University College London Hospitals Biomedical Research Centre. This work was supported by the Medical Research Council [grant number MR/L016311/1]. AE was supported by an award from the International Progressive MS Alliance, award reference number PA-1412-0242.

## Notes

https://github.com/ucl-pond/pySuStaIn

### Competing Interest Statement

The authors have declared no competing interest.

